# Activated CaSo_4_-Induced Vacuolation As A Quantitative Platform For Phagocytosis-Driven Drug Screening

**DOI:** 10.64898/2026.03.11.711143

**Authors:** Vishakha Goswami, Abul Faiz, Gaurav Dutt, Abhishek Kumar, Saima Bashir, Arushi Gupta, Snehasish Das, Alpana Joshi, Subrata K Das

## Abstract

Cytoplasmic vacuolization is a fundamental process associated with phagocytosis, lysosomal acidification, and autophagy, yet robust in-vitro models for its quantification and pharmacological screening remainlimitedor insufficiently established. In this study, we demonstrate that thermally activated calcium sulfate (ACS) induces extensive vacuolation across mammalian cell lines including HeLa, RAW 264.7, 3T3-L1, and SH-SY5Y, thereby establishing a versatile platform to study vacuole biogenesis. To ensure reproducibility, particle heterogeneity was addressed using sedimentation-based fractionation, with homogeneous suspensions obtained at the 5th minute producing stable and consistent vacuole formation. Vacuolation was subsequently quantified by Neutral Red (NR) uptake, dose and time dependent response analyses confirmed direct correlation between ACS concentration and vacuole induction. The assay was validated with bafilomycin A1 (BFA1), a selective V-ATPase inhibitor, which served as a positive control and demonstrated concentration and time dependent inhibition of vacuole formation and acidification. Building on this framework, ten commercially available drugs were screened, revealing distinct profiles ranging from early cytotoxicity, strong vacuole inhibition to partial suppression or negligible effects. This dual capacity to discriminate between vacuole inhibition and cytotoxic responses highlights the utility of ACS-induced vacuolization as a sensitive and scalable in vitro platform. Collectively, our findings position this system as a tractable assay for mechanistic studies of vacuole biology and a functional screening tool for identifying modulators of lysosomal and phagocytic pathways relevant to infection, Lysosomal Disorders, and Phagocytotic dysfunction disorders.

**GRAPHICAL ABSTRACT:** 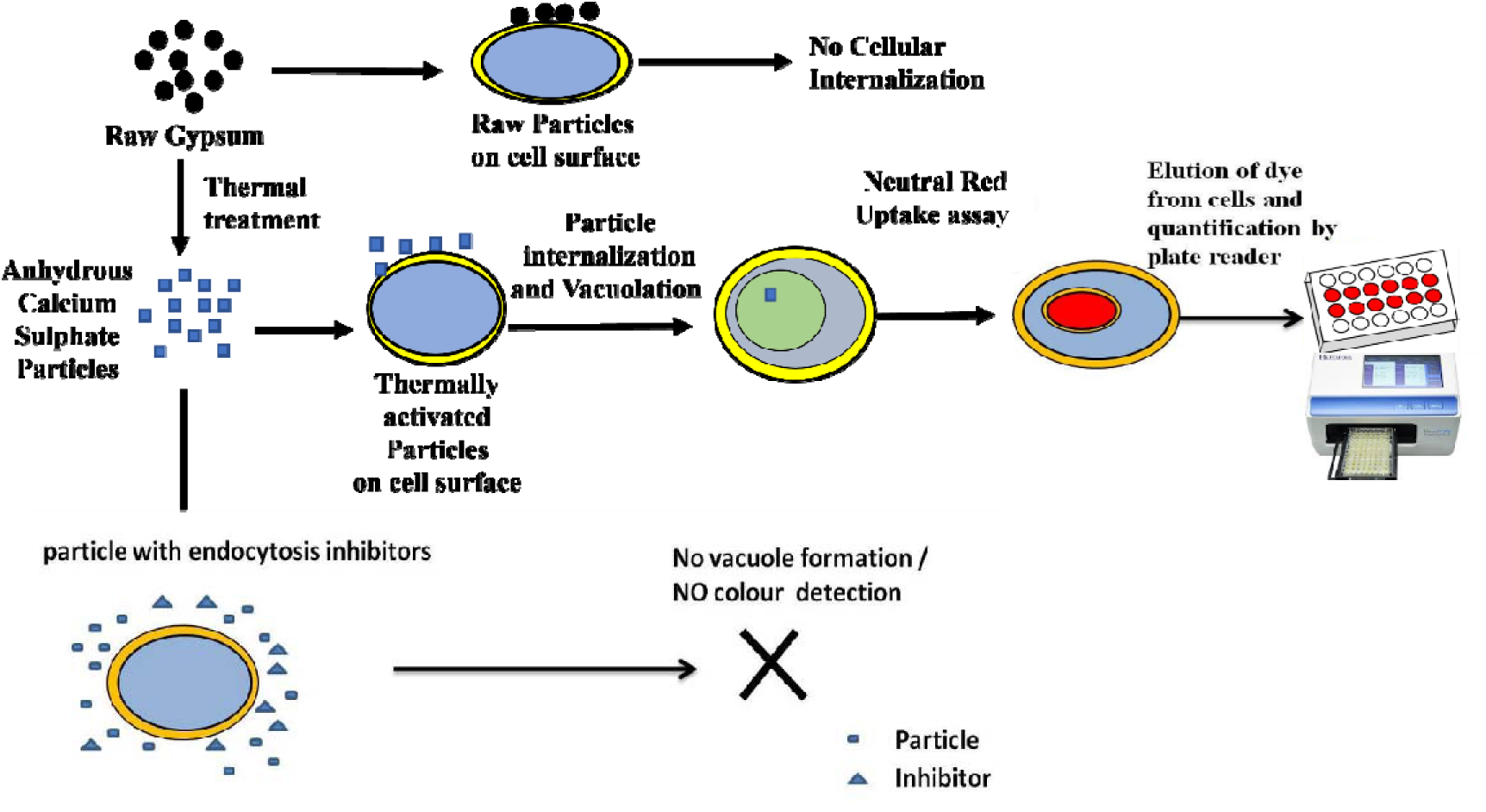

## 1. INTRODUCTION

Cytoplasmic vacuolization, characterized by extensive vacuole formation within animal cells both in vivo and in vitro, can arise spontaneously or be induced by bacterial and viral agents, as well as diverse low-molecular-weight compounds [1–2]. In mammalian cells, vacuolization manifests as either transient or irreversible responses: transient vacuolization occurs exclusively in the presence of an inducing agent and reverses upon its removal [3–4], whereas irreversible vacuolization correlates with cytotoxic states and often culminates in cell death under sustained exposure. Numerous natural and synthetic compounds, including pharmaceuticals and pollutants, have been reported to induce irreversible vacuolization [5–6]. Bacterial toxins and viral proteins frequently act as vacuolating factors in infected cells, with vacuolating proteins serving as key mediators of cytotoxic effects [7–8]. Vacuole biogenesis is closely linked to fundamental cellular processes such as phagocytosis and endocytosis. Vacuoles form when animal cells remodel their plasma membrane to engulf foreign particles (e.g., bacteria, fungiand other exogenous particles) or endogenous materials (e.g., apoptotic cells, infected cells etc) through phagocytosis, which is efficient for particles within specific size ranges [9–10]. Vacuoles may also act as intracellular trafficking vesicles that regulate receptor signalling, recycling, and degradation, collectively categorized under endocytosis. Endocytosis encompasses diverse internalization pathways: smaller particles (<0.5 µm) are typically internalized via receptor-mediated endocytosis or pinocytosis, whereas larger particles are engulfed through phagocytosis [11].

This immune mechanism enables macrophages and other phagocytes(non-professional phagocytes) to internalize pathogens and particles in a ligand-specific manner, often utilizing scavenger receptors for bacteria (0.5–3 µm) and larger particles such as yeast (3–5 µm) [12–13]. Opsonization, in which complement proteins or antibodies coat particles, further enhances recognition and uptake through complement or Fc receptors [14–15].

The initiation of phagocytosis involves specialized receptors recognizing exogenous particles or microbial pathogens, directly or via opsonins [16–17]. Multiple receptor classes can operate simultaneously on a single phagocyte [18]; for instance, Toll-like receptors (TLRs) recognize pathogen-associated molecular patterns (PAMPs), priming the cell for uptake by activating integrins [19–20]. Membrane ruffles and actin-rich lamellipodia form phagocytic cups that seal into vacuoles known as phagosomes or macropinosomes [21–22]. Phagosomes subsequently fuse with lysosomes to form phagolysosomes enriched with hydrolytic enzymes for particle degradation [23]. As they mature through sequential interactions with the endolysosomal system, phagosomes undergo progressive acidification, facilitated by proton-pumping V-ATPase and degradative enzymes [24–25]. The actin cytoskeleton provides the structural force required for engulfing larger particles [26].Pharmacological modulators of these pathways serve as valuable tools for dissecting cellular mechanisms. Bafilomycin A1 (BafA1), a potent inhibitor of V-ATPase, prevents proton pumping across endolysosomal membranes, thereby impairing organelle acidification [27–28].

Phagocytosis and phagosome maturation are essential for immune defense and cellular homeostasis, and their dysregulation contributes to infections, inflammation, cancer, lysosomal disorders and hyper-phagocytic states that are implicated in immune-mediated disorders characterized by pathological engulfment of host cells. Given the clinical need to modulate excessive or pathological phagocytic activity, there is a need for robust, scalable, and cost-effective in vitro phagocytosis assays suitable for pharmacological screening. Conventional assays largely depend on fluorescently labelled particles or synthetic beads, which introduce labelling-associated artifacts, require specialized reagents, and limit throughput. [29-30-31-32-33].

Here, we developed biocompatible activated calcium sulfate (ACS) microparticles that are readily internalized by cells, leading to robust vacuolization. Leveraging this response, we established a neutral red (NR)–based microplate assay to quantitatively measure vacuolization via vacuolar acidification, enabling high-throughput drug screening.

## 1. MATERIAL AND METHODS

### 2.1 Preparation of Activated Calcium Sulphate particles

Raw gypsum (CaSO_4_·2H_2_O) was coarsely powdered and transferred to a ceramic vial. The material was thermally activated by calcination in a muffle furnace at 700 °C for 1 h, followed by passive cooling to room temperature. The resulting activated calcium sulfate (ACS, CaSO_4_) was stored in the borosilicate glassvial under dry conditions for subsequent experiments.

### 2.2 Characterization of Activated Calcium Sulphate particle

Fourier-transform infrared (FT-IR) spectroscopy was performed to assess structural differences between gypsum and activated calcium sulfate (ACS). Spectra were acquired at room temperature using an AGILENT CARY 630 FT-IR spectrometer equipped with an attenuated total reflectance (ATR) module and operated via MicroLab PC software. Measurements were conducted in triplicate using three independent samples per condition to ensure reproducibility.

Powder X-ray diffraction (XRD) analysis was carried out on gypsum and ACS samples using a high-resolution X-ray diffractometer (PANalytical High-Resolution XRD-I, PW 3040/60) equipped with a Cu Kα radiation source (λ = 1.5406 Å). Diffraction patterns were collected over a 2θ range of 10°–80° with a step size of 0.02° and a dwell time of 1 s. The X-ray generator was operated at 40 kV and 30 mA. Phase identification was performed using the JCPDS database, and crystallite size was estimated using the Scherrer equation. Peak fitting and background subtraction were carried out using X’Pert® analysis software.

### 2.3 Preparation of Activated Calcium Sulfate Suspensions and Sedimentation-Based Fractionation

Activated calcium sulfate (ACS) suspensions were prepared by dispersing 100 mg of ACS in 7 mL of Dulbecco’s Modified Eagle Medium (DMEM) supplemented with 10% fetal bovine serum (FBS) in 9 separate sterile 15 mL centrifuge tubes. Each Suspension was vortexed to achieve uniform particle dispersion.

At every 1-min intervals, 5 mL aliquots were carefully aspirated from the upper phase of each suspension and transferred to fresh tubes. This stepwise sedimentation procedure was continued up to 9 min, yielding nine distinct ACS fractions corresponding to sedimentation times of 1–9 min. Each sedimentation fractions was gently pipetted to achieve uniform particle dispersion, and each tube was treated as an independent stock for time-dependent sedimentation analysis. From each fraction, 30 µL of suspension was mounted on glass slides, dried and examined using an inverted microscope (EA PRIME Inverted) to evaluate particle size distribution and the progression from heterogeneity to relative homogeneity (n=3).

### 2.4 Cell Culture and Seeding

Dulbecco’s Modified Eagle Medium (DMEM), fetal bovine serum (FBS), antibiotics, and Trypsin–EDTA were purchased from Thermo Fisher Scientific. HeLa, 3T3-L1, SH-SY5Y, and RAW 264.7 cell lines were obtained from the National Centre for Cell Science (NCCS), Pune, India. Cells were maintained in DMEM supplemented with 10% heat-inactivated FBS, 100 µg/mL penicillin, 100 µg/mL streptomycin, and 5 µg/mL amphotericin B under standard culture conditions (37 °C, 5% CO_2_, and 95% relative humidity).

Stock cultures were grown in 25 cm^2^ tissue culture flasks and sub-cultured using Trypsin–EDTA. Briefly, cells were washed twice with 1x phosphate-buffered saline (PBS), detached with 500 µL Trypsin–EDTA, and neutralized with 1 mL DMEM. The resulting cell suspension was gently pipetted to ensure uniform dissociation, transferred to a 15 mL centrifuge tube, and centrifuged at 1000 rpm for 5 minutes. The supernatant was carefully discarded, and the cell pellet was resuspended in DMEM to a final volume of 10 mL.

For experiments, cells were seeded into 96-well plates by dispensing 90 µL of the cell suspension (10,000 cells) per well and incubated overnight under standard culture conditions to achieve approximately 70% confluency prior to treatment.

### 2.5 Vacuole Induction and Quantification

Vacuole induction experiments were performed 24 h after cell seeding in 96-well plates. For treatment, 30 µL of each activated calcium sulfate (ACS) stock sedimentation fraction (1–9 min) was resuspended by gentle pipetting and added to individual wells containing pre-seeded HeLa cells (n=3). Following overnight incubation under standard culture conditions, cells were examined using a bright-field inverted microscope and subsequently subjected to neutral red (NR) staining for vacuole quantification.

For comparative analysis, the 5-min sedimentation fraction, representing a relatively homogeneous ACS particle population, was similarly resuspended, and 30 µL of the fraction containing smaller particles was added to wells containing HeLa, 3T3-L1, SH-SY5Y, and RAW 264.7 cells (n=3). Cells were incubated overnight and processed for NR staining to assess vacuole formation. ACS-treated cells served as the particle-treated control, while cells cultured without ACS were used as the untreated control.

Quantitative NR uptake data were analyzed, and bar graphs were generated using Microsoft Excel to depict the relationship between ACS particle exposure and vacuole formation.

### 2.6 Dose and Time Dependent Response of Activated Calcium Sulfate Particles

To evaluate dose-dependent cellular responses, the 5-min stock sedimentation fraction of activated calcium sulfate (ACS), was used for the experiment. The particle–medium suspension was resuspended by thorough pipetting prior to dilution to ensure homogeneity.

Serial dilutions of the ACS 5-min sedimentation fraction were prepared to generate graded particle concentrations. For the initial dose range, aliquots of 120 µL, 60 µL, and 30 µL of the small-particle suspension were transferred into separate 1.5 mL microcentrifuge tubes (n = 3 per dose), dried, and weighed independently to determine particle mass. In parallel, corresponding volumes of DMEM (120, 60, and 30 µL) were transferred into separate tubes, dried, and weighed (n = 3) and used as volume-matched control masses for the respective ACS aliquots. It was determined that 120 µL, 60 µL, and 30 µL of the suspension contained approximately 2.5 mg, 1.25 mg, and 0.625 mg, respectively.The corresponding aliquots of 120 µL, 60 µL, and 30 µL from the 5-min sedimentation fraction were added to individual wells of a 96-well plate containing pre-seeded HeLa cells, representing high, medium, and low particle doses, respectively.

To assess vacuolation at lower particle concentrations, further stepwise serial dilutions were performed. Briefly, 30 µL of the small-particle suspension was mixed with 30 µL DMEM, and 30 µL of this mixture was added to a well containing HeLa cells. The remaining suspension was sequentially diluted by the addition of 30 µL DMEM, and 30 µL from each dilution was transferred to successive wells, thereby generating progressively lower particle concentrations.Following ACS exposure, cells were incubated overnight under standard culture conditions (n=3). Vacuole formation was assessed by neutral red (NR) staining, and stained cells were examined by microscopy. NR uptake was quantified, and the resulting data were analyzed and presented as bar graphs to depict the relationship between ACS particle dose and vacuolation.

To evaluate time-dependent responses, the 5-min stock sedimentation fraction of ACS was used. The particle–medium suspension was resuspended and subsequently, 30 µL of the fraction containing smaller particles was added to the wells of 96-well plate pre-seeded with HeLa cells. Neutral Red (NR) staining was performed to assess vacuolation at defined time points. Briefly, 60 µL of Neutral Red solution was added at 0, 6, 12, 24, 36, 48, 60, 72, and 84 h post-treatment (n = 3 per time point)and the resulting datawere analyzed.

### For comparative analysis, time-lapse imaging was performed to assess time-dependent cellular responses in A549 cells treated with 2-min stock-sedimented ACS particles.2.7 Dose-Dependent Vacuole inhibition using Bafilomycin A1

Bafilomycin A1 (100 µg; MedChemExpress) was used to evaluate dose-dependent inhibition of vacuole formation. Stock solution of Bafilomycin A1 (BFA1) at a concentration of 100 µg/mL (160.5 µM) was prepared by dissolving 100 µg of BFA1 in 50 µL of DMSO, followed by dilution in 950 µL of DMEM. 1 µL of the stock solution was diluted in 1.6 mL of DMEM to obtain a working solution of 100nM. The solution was mixed thoroughly by repeated pipetting to ensure uniform distribution.

Dose-dependent distribution of Bafilomycin A1 was performed by adding 95 µL to 5 µL of the 1.6 mL working solution to wells of a 96-well platepre-seeded with HeLa cells. Following Bafilomycin A1 exposure, 30uL of5-min stock sedimentation fraction of ACS suspension was added to each well. Cells were then incubated overnight under standard culture conditions to induce vacuole formation(n=3).

After incubation, cells were stained with neutral red (NR) and examined to determine the concentration of BFA1 at which vacuole formation was first observed. NR uptake by vacuoles was quantified, and the data were analyzed. A bar graph was generated using Microsoft Excel to illustrate the correlation between Bafilomycin A1 concentration and vacuole formation.

### 2.8 Delayed inhibition of vacuole acidification by Bafilomycin A1

To evaluate the effect of BFA1 on vacuole dynamics, HeLa cells in 96-well plates were first induced to form massive vacuoles by the addition of 30ul of 5^th^ minute ACSstock particles and incubated overnight, under standard temperature and conditions. Following vacuole formation, 95 µL (51.35nM) of BFA1 working solution was administered under three distinct experimental conditions to evaluate its time-dependent effects on vacuolar acidification, which was assessed by performing Neutral Red (NR) staining at multiple time points via microscopy. For immediate responses, NR staining was conducted at 0, 15, 30, 45, 60, and 75 minutes following BFA1 addition. For intermediate responses, BFA1 was introduced into cells containing pre-formed vacuoles, and NR staining was carried out at 0 min and at hourly intervals up to 6 hours. For long-term responses, BFA1 was added to cells with established vacuoles, and NR staining was performed daily from Day 0 to Day 2. In all cases, ACS-treated cells without BFA1 were maintained ascontrols and stained with NR at the corresponding time points.

### 2.9 Neutral Red staining

Relative vacuolation was measured in mammalian cells using the Neutral Red dye [33–34]. The experiments took place in 96-well plates. Neutral red (0.5 mg/ml) was added to each well, incubated for 30 minutes in a CO_2_ incubator, washed two times with 1x PBS, and eluted with a destining solution (100 µl of 50% dehydrated ethanol, 49% deionized water, and 1% glacial acetic acid). Neutral Red absorption was monitored by measuring the absorbance at 492 nmwith Elisa plate reader (Model:ER-181s).

### 2.10 Acridine Orange Uptake Assay

3T3-L1 Cells were cultured on cavity slides by adding 100 µL of cell suspension to two cavities per slide, leaving the middle cavity empty to prevent cross-contamination. Each cavity was supplemented with 50 µL of Dulbecco’s Modified Eagle Medium (DMEM) and incubated at 37°C in a humidified atmosphere containing 5% CO_2_ for 24 hours to facilitate cell adherence and stabilization.

Cells were treated with5-minute sedimentation fraction ofACS to induce vacuole formation and incubated for an additional 24 hours under standard experimental conditions. To evaluate the effect of bafilomycin A1 on vacuolar dynamics, 95 µL of bafilomycin A1 working solution was added to one cavity, while the second cavity served as an untreated control. Both cavities were incubated under the same conditions for another 24 hours.

Following incubation, cells were stained with acridine orange solution (1mg/mL) to selectively label vacuoles and acidic vesicular organelles. The staining was performed by incubating the cells with the dye for 15 minutes at 37°C. Post-staining, the cells were washed thrice with sterile phosphate-buffered saline (PBS) to remove excess dye and reduce nonspecific background fluorescence.

The stained slides were immediately examined using a fluorescence microscope(Model: DiGi 110 Infinity) to assess cellular morphology and acridine orange uptake, enabling the characterization of vacuolar acidification and dynamics.

### 2.11 Drug Screening

An in-vitro drug screening assay was developed to identify inhibitors of phagocytosis-associated vacuole formation. Bafilomycin A1–based vacuolar inhibition was usedas a positive control. Commercially available pharmaceutical formulations were used for drug preparation. For solid formulations, tablets containing ≤5 mg of active ingredient were processed using two tablets, whereas tablets containing >5 mg of active ingredient were processed using a single tablet. Tablets were transferred to sterile 15 mL centrifuge tubes, resuspended in 7 mL sterile phosphate-buffered saline (PBS), and vortexed until complete dispersion or solubilization. The suspensions were centrifuged at 3000 rpm for 10 min to pellet insoluble excipients, and the clarified supernatant was collected and stored at 4°C until further use. Liquid drug formulations were used directly, and 100 µL of the undiluted preparation was taken as the stock solution.

For drug screening experiments, columns 1–10 of a 96-well plate were used to evaluate 10 different drugs, with each compound serially diluted across wells within its respective column. Column 11 served as the drug positive control; the first four wells received 51.35, 48.57, 45.45, and 41.9nM of BFA1 working solution, respectively, while the remaining four wells received 28, 21.74, 14.29, and 5.26nMof the BFA1 working solution, in accordance with the dose-dependent vacuole inhibition protocol described above.

Column 12 served as the control column, with the first four wells receiving 5-min sedimentation fraction of ACS suspension alone (without drug treatment) and the remaining wells containing untreated cells without ACS or drugs.

Serial dilutions of each drug stock were prepared in DMEM to generate a concentration gradient. Briefly, 100 µL of the drug stock solution was mixed with 100 µL DMEM, and 100 µL of the resulting dilution was added to the first well of a 96-well plate pre-seeded with HeLa cells. Subsequent dilutions were generated by sequential addition of 100 µL DMEM to the remaining solution, thorough mixing, and transfer of 100 µL to successive wells. This process was repeated across eight wells to obtain eight drug concentrations, with a final volume of 100 µL per well.

Following drug treatment, vacuole formation was induced by the addition of 30 µL of the fifth-minute homogenized ACS particle suspension to each well. Plates were incubated at 37 °C in a humidified atmosphere containing 5% CO_2_ for 24 hrs.Neutral red (NR) dye (60 µL) was then added directly to each well and incubated for 15 min at 37 °C. Cells were washed twice with PBS to remove excess, non-internalized dye. Vacuolar NR uptake was visualized using a bright-field inverted microscope and quantified using a microplate reader. Quantitative data were analyzed, and dose-dependent effects of drug treatment on vacuolar NR uptake were represented graphically.

## 3. RESULT AND DISSUCATION

### 3.1 Characterization of anhydrous ACS via FT-IR and XRD

Thermal activation of calcium sulfate induced a clear dehydration-driven phase transition from gypsum (CaSO_4_·2H_2_O) to anhydrous activated calcium sulfate (CaSO_4_). FT-IR analysis provided direct chemical evidenceof this transformation. Raw gypsum exhibited prominent O–H stretching and bending vibrations in the regions of 3000–4000 cm^−1^ and 1600–1700 cm^−1^, respectively, corresponding to hydrogen-bonded water molecules within the gypsum lattice (Figure 1A). In contrast, heat-activated calcium sulfate (ACS) showed complete disappearance of these O–H vibrational bands (Figure 1B), confirming the removal of crystalline water and formation of an anhydrous phase. This disappearance of O-H vibrational bands upon heating is a definitive indication of the removal of crystalline water molecules [35].Characteristic sulfate (SO₄^2^⁻) vibrational bands observed around 1100–1150 cm^−1^ were retained in both samples, with minor shifts in the activated material. The overall changes indicated the internal structural rearrangement and how it behaves physically and chemically [36].

**Figure 1.**
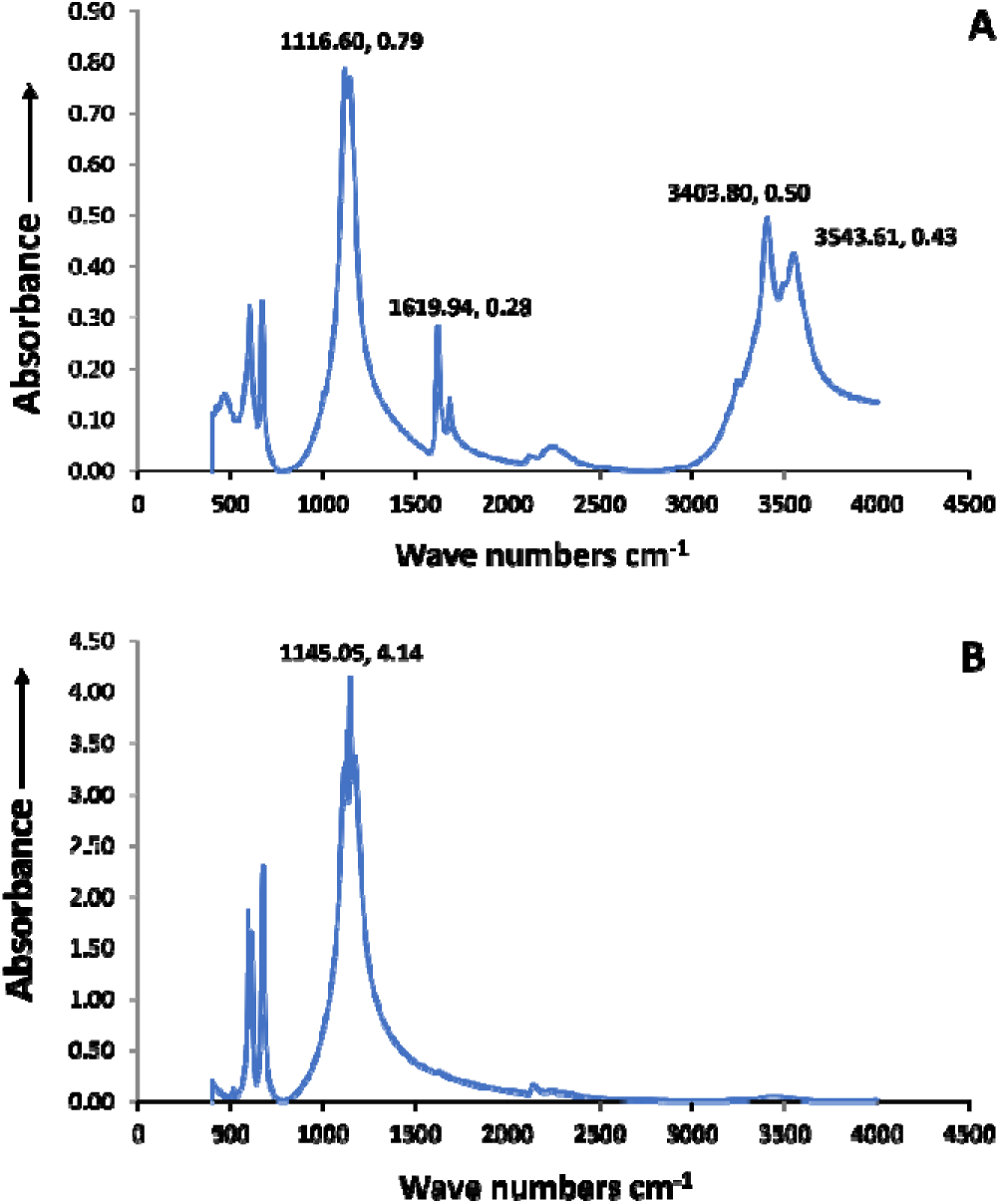
FTIR analysis of raw and heat-activated calcium sulfate. (A) FTIR spectrum of calcium sulfate dihydrate (CaSO_4_·2H_2_O) showing characteristic O–H stretching (3000–4000 cm^−1^) and bending (1600–1700 cm^−1^) vibrations corresponding to crystalline water, along with sulfate group (SO₄^2^⁻) absorption bands around 1100–1150 cm^−1^. (B) FTIR spectrum of heat-activated calcium sulfate (CaSO_4_), demonstrating the disappearance of O–H vibrational bands, confirming the removal of crystalline water and transition to the anhydrous phase.

**Figure 2.**
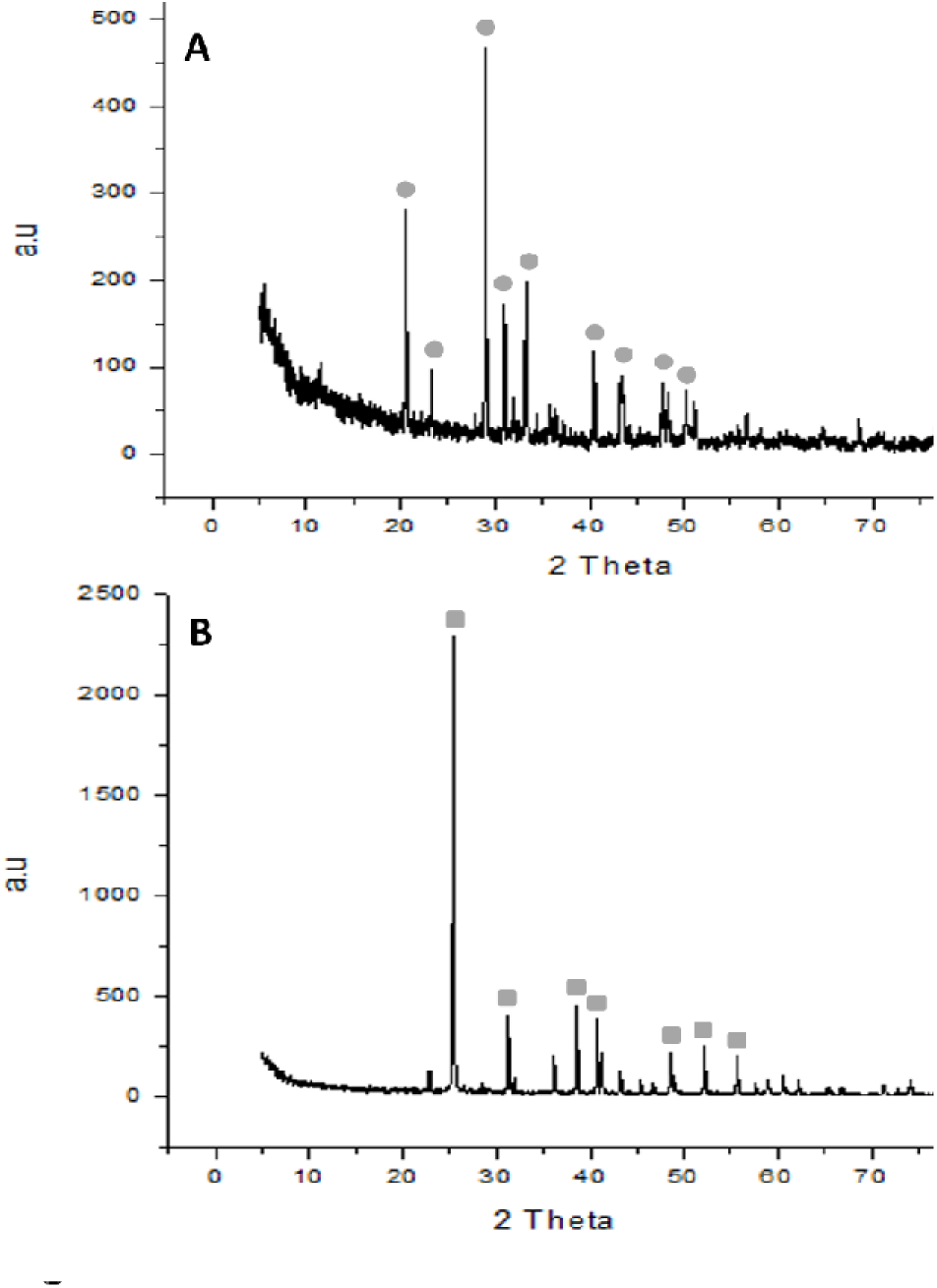
XRD patterns of hydrated and anhydrous calcium sulfate. (A) XRD profile of calcium sulfate dihydrate (CaSO_4_·2H_2_O) exhibiting diffraction peaks at 2θ values of 20.5°, 23.2°, 29°, 31°, 33.2°, 40.4°, 43.4°, 48°, and 50°, characteristic of the crystalline gypsum structure. (B) XRD profile of heat-activated anhydrous calcium sulfate (CaSO_4_), displaying distinct peaks at 2θ values of 25.2°, 31°, 36°, 38.2°, 40.9°, 48.4°, 52°, and 55.9°, confirming structural transformation to the anhydrite phase.

**Table 1.** List of drugs used for vacuole modulation screening in ACS-treated HeLa cells. A panel of 11 pharmacological agents with diverse therapeutic classes was selected for screening. The table details the drug name and the administered concentration, expressed either as milligram content or percentage composition, as appropriate. These compounds were systematically tested for their ability to modulate vacuole formation and neutral red uptake in HeLa cells following treatment with activated calcium sulfate (ACS).

| Sl No | Drugs | Mg |
| --- | --- | --- |
| 1. | Ecosprin-75 | 75mg |
| 2. | Zerodol-SP | 440mg |
| 3. | Galpride M1 | 500mg |
| 4. | Zycel | 200mg |
| 5. | Atorlip | 10mg |
| 6. | Otrivin Oxy | 0.5% |
| 7. | Pred Forte | 1% |
| 8. | Tropag-P | Phenylephrine (5%) +<br>Tropicamide (0.8%) |
| 9. | O2 Tablets | 700mg |
| 10. | BE-Fame-SP | 440mg |

X-ray diffraction (XRD) serves as an indispensable tool for characterizing the crystalline phases of materials and monitoring phase transitions.The XRD spectrum was analyzed for both raw calcium sulfate dihydrate (CaSO_4_·2H_2_O) particles and heat-activated anhydrous calcium sulfate particles. For the dihydrated calcium sulfate, characteristic diffraction peaks were observed at 2θ values of 20.5°, 23.2°, 29°, 31°, 33.2°, 40.4°, 43.4°, 48°, and 50°, corresponding to the well-defined crystalline structure of gypsum (Figure-2A). These peaks arise due to the ordered arrangement of calcium, sulfate, and water molecules in the lattice. In contrast, the heat-activated anhydrous calcium sulfate displayed diffraction peaks at 2θ values of 25.2°, 31°, 36°, 38.2°, 40.9°, 48.4°, 52°, and 55.9°, indicating a structural transformation into an anhydrous phase (Figure-2B). The diffraction peaks observed for gypsum correspond to its monoclinic crystal system, with water molecules playing a pivotal role in maintaining lattice stability and symmetry [37-38-39]. Upon heating, the dehydration process results in significant structural rearrangements, giving rise to new peaks characteristic of anhydrite. The shift in peak positions and the emergence of new peaks in the anhydrous form signify a reduction in crystal symmetry, attributed to the collapse of the hydration framework [37-38-39-40].

### 3.2 Particle Heterogeneity/Homogeneity in Vacuole Biogenesis and Its Quantification

Particle heterogeneity is a critical determinant of cellular uptake dynamics and vacuolar responses, as variations in particle size, shape, and aggregation state directly influence phagocytic efficiency and intracellular processing [41]. In our previous work, we demonstrated thatGodanti Bhasma (formulation of anhydrous CaSO4 with herbal extract) induces massive vacuolation in mammalian cells, whereas gypsum fails to elicit comparable cellular responses [42]. In the present study, raw gypsum was coarsely powdered and thermally activated to generate activated calcium sulfate (ACS) microparticles capable of inducing pronounced vacuolation in mammalian cells. However, the vacuolating activity of ACS was heterogeneous, likely reflecting variability in particle characteristics. Owing to the mechanical nature of the preparation process, the resulting particle suspensions initially comprised a broad distribution of particle sizes.

Microscopic examination was performed on ACS suspensions collected at each sedimentation time point, while neutral red (NR) staining was carried out on HeLa cells treated with ACS derived from the corresponding fractions. Early sedimentation fractions (1–3 minutes) contained highly heterogeneous particle populations, characterized by wide size variability and elevated particle numbers (Supplementary Figure-1F-1H). HeLa cells exposed to these early fractions exhibited extensive vacuolation, consistent with robust phagocytic uptake of particles spanning a broad size range (Supplementary Figure-1A-1C). From the 4-minute time point onward, ACS preparations showed a progressive shift toward homogeneity, with particles displaying a more uniform size distribution. By the 5-minute fraction, particle numbers stabilized, and suspensions remained consistent across subsequent time points (Supplementary Figure-1I-1J). Correspondingly, NR staining revealed that vacuolation in HeLa cells treated with ACS from the 5–9-minute fractions was moderate, uniform, and highly reproducible (Figure 3).This sedimentation-based separation thus provided a temporal means of enriching for particles of different size classes. At early time points (1–3 minutes), incomplete sedimentation resulted in suspensions enriched in larger and more polydisperse particles. Such heterogeneity likely enhanced phagocytic activity, as cells encounter both larger particles that require extensive actin remodelling and smaller particles that are rapidly internalized (Figure-3). The mixed particle load could trigger multiple uptake pathways simultaneously, producing pronounced vacuolation. In contrast, later fractions—particularly the 5-min fraction—predominantly contain smaller, sedimentation-stable particles that produce a uniform uptake profile and consistent vacuolar responses. Based on these criteria, the 5-min sedimentation fraction was selected for all subsequent cellular experiments to ensure reproducibility and quantitative consistency.

**Figure 3.**
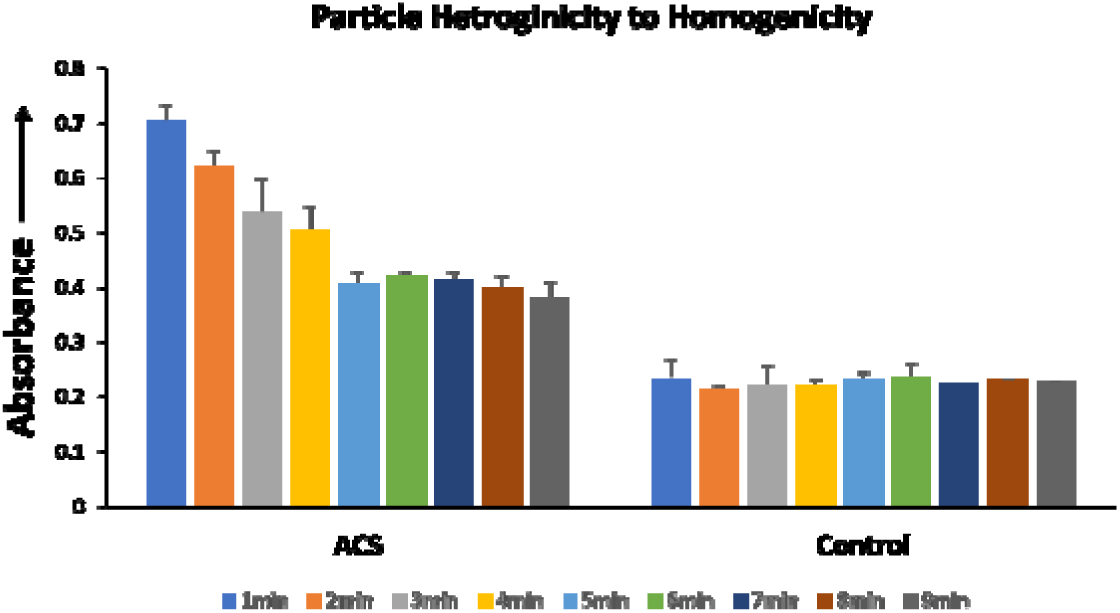
Correlation of particle homogeneity with reproducible vacuolation. HeLa cells exposed to early heterogeneous suspensions (1–3 minutes) exhibited pronounced and variable vacuolation, while later homogeneous suspensions (5–9 minutes) produced more consistent and moderate vacuolation patterns. These results demonstrate the direct impact of particle heterogeneity on cellular uptake dynamics and vacuole formation (n=3).

This method was established to quantify vacuolation across different cell lines, in HeLa, 3T3-L1, RAW-264.7, and SH-SY5Y cells (Figure 4B–3I). Untreated cells contained only small lysosomal vacuoles, whereas 5^th^ minute ACS stock-treated cells displayedabundant vacuoles that were stained with neutral red (NR), confirming their acidic nature [43]. Since lysosomal and vacuolar compartments maintain a lower pH than the cytoplasm, NR readily penetrates and accumulates in these organelles. Quantitative bar graph analysis of NR uptake further supported ACS-induced vacuolization (Figure 4A).In contrast to conventional phagocytic substrates such as latex beads or zymosan, ACS consistently induces extensive vacuolation characterized by large, optically clear, and stable phagosomes that are readily quantifiable [42–43]. These oversized vacuoles constitute a pronounced and reproducible cellular phenotype, permitting high-resolution observation of vacuole formation andacidification. Consequently, ACS provides a uniquely interpretable in vitro phagocytic model that is not attainable using commercially available particles. Furthermore, ACS exhibits strong compatibility with Neutral Red (NR) staining, wherein the formation of large vacuoles facilitates a simple and cost-effective assessment of vacuolar acidification. Consistent with this, NR robustly accumulated within ACS-induced vacuoles [42].

**Figure 4.**
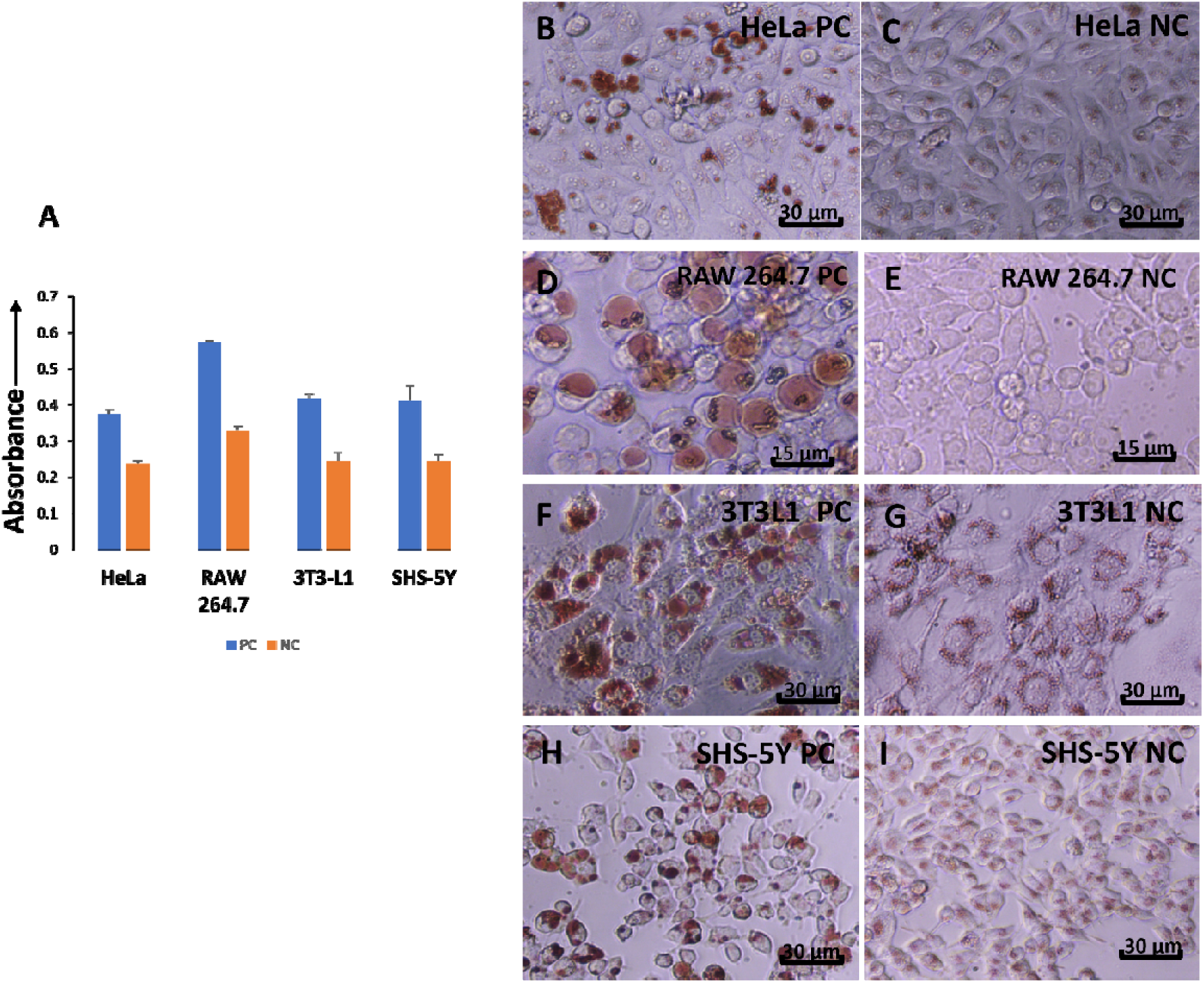
Quantification of ACS-induced vacuolation in mammalian cells. (A) Bar graph analysis of neutral red (NR) uptake demonstrating significantly elevated vacuolation in 5-minsedimentation ACS stock -treated cells compared to controls. (B–I) Representative microscopy images of HeLa, 3T3-L1, RAW-264.7 (Scale= 15µm), and SH-SY5Y cells showing abundant ACS-induced vacuoles stained with NR, confirming their acidic nature, in contrast to untreated controls that contained only small lysosomal vacuoles (n=3) (Scale=30µm).

### 3.3 Concentration and Time-Dependent Cellular Effects of Activated Calcium Sulfate Particles

Quantification of ACS-induced vacuolation was performed using the neutral red (NR) uptake assay in HeLa cells treated with the 5-minute ACS sedimentation fraction. A clear dose-dependent response was observed, with vacuole formation increasing progressively with rising concentrations of ACS particles. Vacuole formation and acidification were assessed using Neutral Red staining, which preferentially labels acidic intracellular compartments.

At the highest concentration tested (120 µL, ∼2.5 mg), HeLa cells exhibited extensive vacuolation, characterized by the formation of numerous vacuoles per cell. This phenotype was readily visualized by light microscopy and was accompanied by robust neutral red (NR) uptake, indicating active acidification of the vacuolar compartments (Figure 5B). At intermediate (60 µL, ∼1.25 mg) and lower (30 µL, ∼0.625 mg) concentrations, vacuole formation was well defined with an optimal and controlled distribution of vacuoles per cell, corresponding to diminished NR accumulation (Figure- 5C-5D).

**Figure 5.**
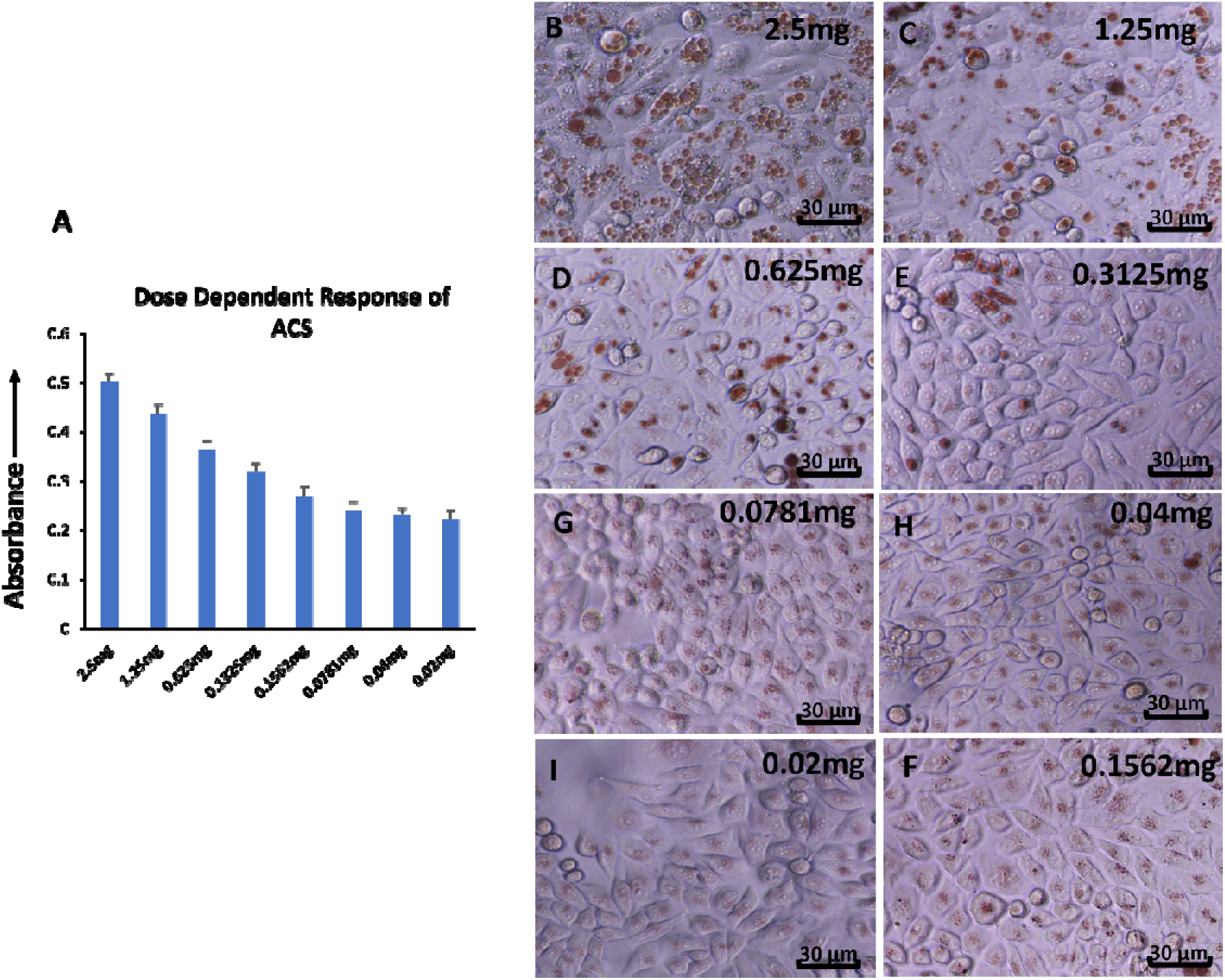
Dose-dependent vacuolation response in ACS-treated HeLa cells. (A) Bar graph analysis of neutral red (NR) uptake showing a positive correlation between ACS concentration and vacuole formation. (B-D) Microscopy images of HeLa cells treated with 120 µL (2.5 mg) ACS exhibiting abundant massive vacuoles, compared with 60 µL (1.25 mg) and 30 µL (0.625 mg) treatments, which produced progressively fewer and smaller vacuoles. (E-I) Serial dilution treatment demonstrating sparse, isolated vacuoles at the lowest concentration, indicating reduced vacuolization. Quantitative comparison across concentrations confirming the threshold-dependent induction of vacuoles (n=3) (Scale= 30µm).

**Figure 6.**
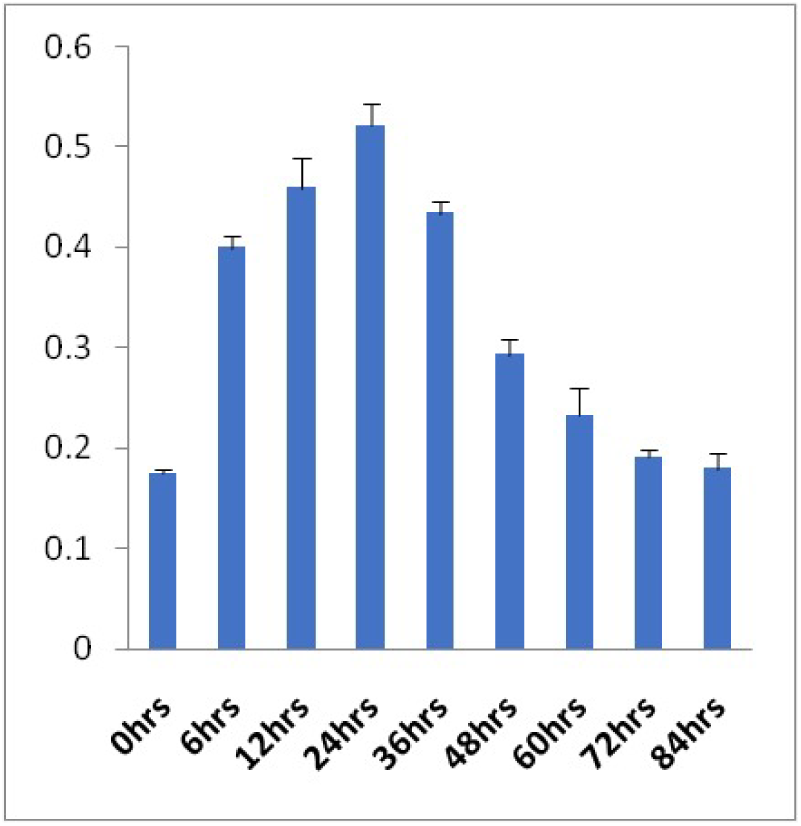
Time-dependent quantification of vacuole formation and turnover in ACS-treated cells using Neutral Red uptake assay. Neutral Red (NR) uptake was measured at indicated time points following treatment with activated calcium sulfate (ACS) particles to assess vacuole formation and intracellular processing. At 0 h, NR absorbance was minimal, indicating basal dye accumulation and absence of ACS-induced vacuoles. NR uptake increased at 6 h, reflecting early vacuole formation during ACS internalization, and further increased at 12 h, reaching a maximum at 24 h, corresponding to extensive formation of large acidic vacuoles. Subsequently, NR absorbance progressively declined from 36 h to 84 h, indicating degradation of internalized ACS particles and vacuolar turnover. NR levels at later time points (48–84 h) were lower than those observed at 6 h, consistent with active vacuolar resolution and restoration of cellular homeostasis. Data represent quantitative measurement of NR absorbance as an indicator of vacuolar dynamics following ACS treatment (n=3).

**Figure 7.**
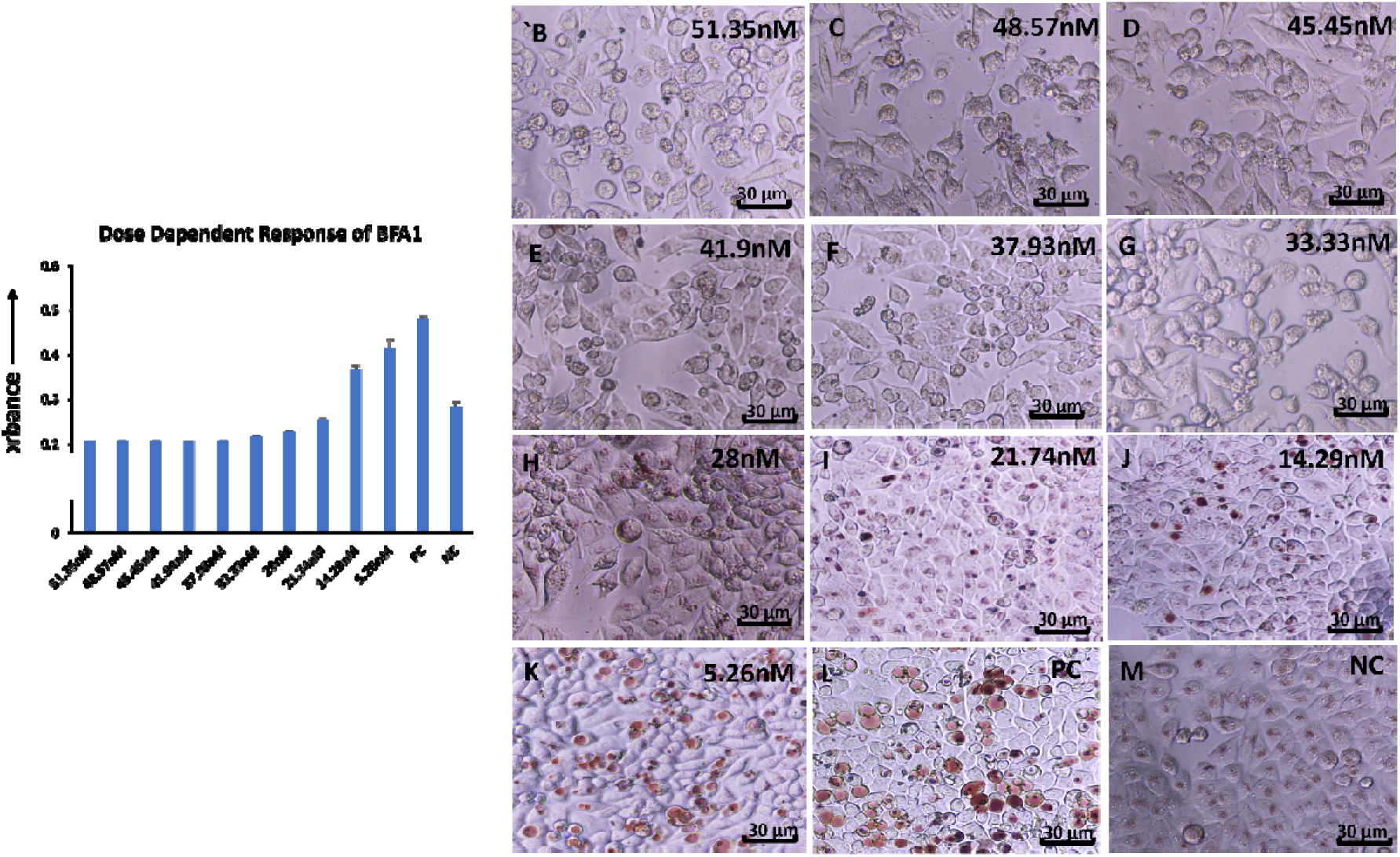
Dose-dependent effects of Bafilomycin A1 on ACS-induced vacuolation and acidification. (A) Bar graph of neutral red (NR) uptake showing the correlation between Bafilomycin A1 concentration and vacuole acidification. (D) Microscopy images of cells treated with 25 µL Bafilomycin A1 displaying small NR-positive vacuoles, while lower concentrations (15 µL and 5 µL) induced abundant large vacuoles with strong NR staining. (C–D) Higher concentrations of Bafilomycin A1 markedly inhibited vacuole formation, consistent with V-ATPase inhibition and loss of phagosome acidification (n=3) (Scale= 30µm).

Further reduction in particle concentration through serial dilution resulted in a pronounced attenuation of vacuolar responses. At the lowest dilution, only isolated vacuoleswere detectable, reflecting minimal vacuole induction relative to higher ACS doses (Figure-5E-5I). Quantitative analysis of NR uptake, presented as a bar graph, corroborated these observations and demonstrated a strong positive correlation between ACS particle concentration and vacuole formation (Figure 5A).

Collectively, these findings establish particle concentration as a critical determinant of ACS-induced vacuolation. Higher particle loads promote robust vacuole formation and acidification, consistent with efficient phagocytic uptake and phagosome maturation, as evidenced by enhanced NR accumulation. In contrast, lower particle concentrations appear insufficient to trigger a sustained phagocytic response, resulting in limited vacuole formation. The serial dilution experiment further supports the existence of a threshold particle concentration required for effective vacuole induction. These results highlight the concentration-dependent regulation of vacuolar dynamics in response to ACS.

Time-dependent cellular responses to activated calcium sulfate (ACS) particles were assessed by quantifying Neutral Red (NR) uptake as an indicator of vacuole formation and dynamics. At 0 h, NR absorbance was minimal, reflecting basal dye accumulation and the absence of ACS-induced vacuoles immediately after particle addition (Figure-6). By 6 h, NR uptake increased, indicating initiation of vacuole formation during early ACS internalization. NR absorbance further increased at 12 h and peaked at 24 h (Figure-6), corresponding to extensive formation of large acidic cytoplasmic vacuoles that actively accumulated NR, consistent with maximal vacuolar expansion and intracellular processing of ACS particles.

Following this peak, NR absorbance progressively declined from 36 h to 84 h, indicating degradation of internalized ACS particles within acidic vacuolar compartments and associated vacuolar turnover and recycling [42]. Notably, NR levels at 48, 60, 72, and 84 h were lower than those at 6 h (Figure-6), supporting active vacuolar resolution and restoration of cellular homeostasis. Time-lapse video of A549 cells treated with 2-min stock sedimentation heterogeneous ACS particles demonstrated a progressive increase in vacuole number and size up to 24 h. Beyond 24 h, both vacuole number and size gradually declined. By day 4, a high degree of vacuolar turnover was observed, although a few isolated vacuoles remained visible (Supplementary Video-1). This persistence is likely attributable to the heterogeneous nature of the 2-min sedimentation fraction, which contains larger ACS particles that require extended time for complete intracellular degradation. Collectively, these findings demonstrate that ACS-induced vacuolation is a dynamic and reversible process involving distinct phases of vacuole biogenesis, maturation, and recycling, which can be quantitatively monitored by NR uptake [42].

### 3.4 Suppression of ACS-Induced Vacuolation by Bafilomycin A1

The effect of bafilomycin A1 on vacuole formation was evaluated by varying its concentration in cultured cells exposed to ACS (activated calcium sulfate). Bafilomycin A1, known for inhibiting the vacuolar-type H+-ATPase (V-ATPase)[44], was used to investigate its role in vacuole formation and acidification. At higher concentrations of bafilomycin A1(51.35nM to 28nM), vacuole formation was significantly inhibited (Figure-7B-7H). When cells were treated with 21.74nM of bafilomycin A1, small vacuoles began to appear, which successfully took up Neutral Red (NR) stain (Figure-7I), indicating the presence of acidic compartments. However, at lower concentrations (14.29 and 5.26nM), the vacuoles increased dramatically in both size and number (Figure-7J&K). These larger vacuoles exhibited active NR uptake (presented as bar graphs, Figure-7A), suggesting that vacuoles formed under these conditions retained their acidic nature and likely underwent normal phagosome maturation processes.

Cell viability remained at approximately 90% even at higher Bafilomycin A1 concentrations, indicating that the concentrations used were not cytotoxic. The data suggest that bafilomycin A1 influences vacuole size and acidification in a dose-dependent manner, with lower concentrations leading to more prominent vacuole formation and acidification.

Bafilomycin A1 has a well-established role as a V-ATPase inhibitor, and its effect on vacuole acidification and maturation is well-documented in various cell types [44]. In this study, we observed that Bafilomycin A1 inhibits vacuole formation at higher concentrations, consistent with its known inhibitory action on proton pumps, which are essential for maintaining the acidic environment within phagosomes. This acidification is critical for phagosome maturation, leading to the recruitment of various hydrolases that aid in particle degradation [45]. The formation of small vacuoles at concentration (21.74nM) of Bafilomycin A1, and their ability to take up NR, implies that V-ATPase activity was still functional, allowing partial acidification of these compartments. Interestingly, at lower concentrations (14.29nM and below), the vacuoles not only increased in number but also in size, all actively taking up NR stain. This suggests that low doses of Bafilomycin A1 may permit vacuole formation and maturation, possibly through residual V-ATPase activity or alternative mechanisms that compensate for the partial inhibition of proton pumps.

These findings highlight the dose-dependent effects of bafilomycin A1 on phagosome dynamics and suggest that low levels of V-ATPase inhibition do not fully block vacuole formation and maturation. Furthermore, the drastic increase in vacuole size at lower concentrations points which may lead tovacuole fusion or vesicle trafficking, which may be upregulated in the absence of full V-ATPase inhibition. This insight could be useful for understanding the fine regulation of vacuole acidification and quantification and its impact on phagocytosis, particularly in macrophage dysfunction models where vacuole formation is impaired or exaggerated [44].

### 3.5 Delayed inhibition of Vacuolar Acidity Following Bafilomycin A1 Treatment

To evaluate the temporal effects of bafilomycin A1 on vacuole dynamics, HeLa cells with pre-formed vacuoles were treated with bafilomycin A1 under three different conditions: immediate, hourly, and daily response assessments. These treatments were followed by Neutral Red (NR) staining to measure vacuole acidification over time.

Immediate Response Analysis: Upon the addition of51.35nM of bafilomycin A1working solution, vacuoles induced by ACS (activated calcium sulphate) particles continued to actively take up NR stain at all-time points observed (0 min to 90 min). This suggests that bafilomycin A1 did not immediately affect vacuole acidification within the first 90 minutes of treatment (Supplementary Figure-2).

Hourly Response Analysis: Over a 6-hour observation period, vacuoles consistently took up NR stain at all hourly intervals (0 min, 1 hr, 2 hrs, 3 hrs, 4 hrs, 5 hrs, and 6 hrs) following Bafilomycin A1 treatment. This indicates that Bafilomycin A1’s impact on vacuole acidification was not significant during the first 6 hours of exposure (Supplementary Figure-3).

Daily Response Analysis: At daily intervals, vacuoles took up NR stain on Day 0 following the addition of bafilomycin A1, similar to the immediate and hourly response analyses. However, after 24 hours of bafilomycin A1 incubation (Day 1), vacuoles no longer took up NR stain. This absence of NR uptake persisted on Day 2, indicating that bafilomycin A1 effectively inhibited vacuole acidification over prolonged exposure (Supplementary Figure-4).

These results suggest a delayed inhibitory effect of bafilomycin A1 on vacuole acidification, as vacuoles remained acidic and functional during the first several hours of treatment but progressively lost their ability to take up NR over extended incubation times.

In the immediate response analysis, vacuoles induced by activated calcium sulfate particles continued to take up NR stain during the first 90 minutes following Bafilomycin A1 treatment, suggesting that vacuole acidification is not immediately halted by V-ATPase inhibition. This indicates a lag phase during which the residual proton gradient and pre-existing acidic conditions are maintained, even in the presence of bafilomycin A1. However, in the daily response analysis, vacuoles lost their ability to take up NR stain after 24 hours of bafilomycin A1 incubation. In the time-dependent response analysis, vacuoles lost their ability to take up neutral red (NR) after 24 hours of bafilomycin A1 treatment, and this loss of acidification persisted at 48 hours, indicating sustained impairment of vacuole maturation. This effect is consistent with inhibition of vacuolar H⁺-ATPase (V-ATPase) activity. The delayed onset likely reflects residual intravacuolar acidity at early time points, which is progressively depleted during ACS degradation and lysosomal enzyme activity, ultimately leading to dissipation of the proton gradient required for vacuolar acidification.The inhibition of vacuole acidification over extended periods highlights bafilomycin A1’s significant role in controlling vacuolar dynamics and phagosome maturation[44].

Overall, these findings underscore the importance of proton pump regulation in phagosome acidification and provide insights into the temporal dynamics of vacuole maturation when V-ATPase is inhibited. Future studies could explore whether other compensatory mechanisms or ion exchangers contribute to the observed delayed inhibition of vacuole acidification by Bafilomycin A1 [44].

### 3.6 Visualization of Acidic Vacuolar Compartments by Acridine Orange

Control cells treated with only ACS and stained with acridine orange displayed prominent orange–red fluorescence within vacuoles, indicating the presence of an acidic intravacuolar environment, while the nuclei fluoresced green, consistent with the differential accumulation of acridine orange in acidic compartments and nucleic acids (Figure 8B). In contrast, ACS-treated cells subsequently exposed to bafilomycin A1 (after 24hrs) exhibited vacuoles that failed to show orange–red fluorescence and remained colourless under fluorescence microscopy, although nuclear green fluorescence was preserved (Figure 8 A). The absence of vacuolar staining indicates a loss of vacuolar acidification following bafilomycin A1 treatment.

**Figure 8.**
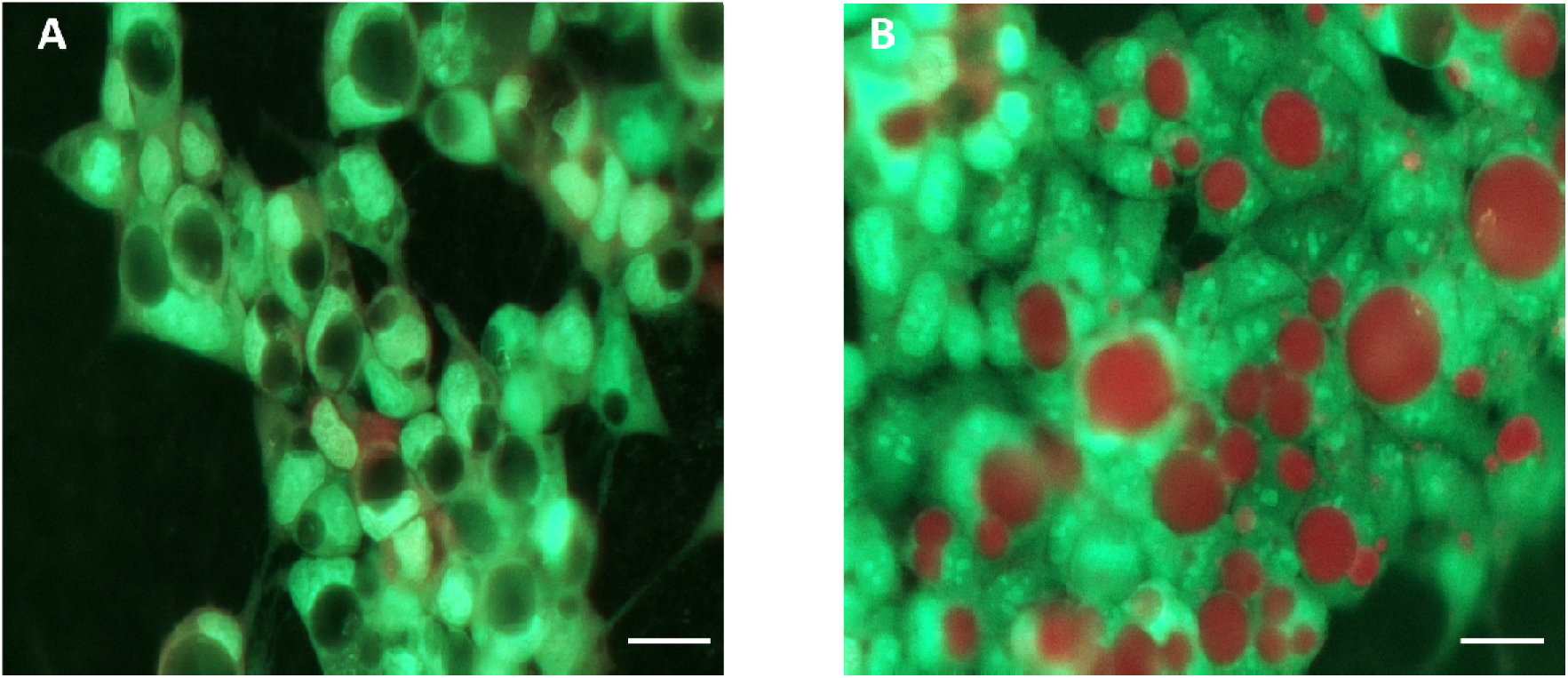
Acridine orange assay demonstrating inhibition of vacuolar acidification by Bafilomycin A1. (A) Fluorescence microscopy images of ACS-treated cells exposed to Bafilomycin A1 showing vacuoles that failed to accumulate acridine orange, appearing colourless, while nuclei fluoresced green. (B) Control cells exhibited strong orange-red fluorescence within vacuoles, confirming acidic compartments, along with green nuclear staining. These results indicate that Bafilomycin A1 selectively blocks vacuolar acidification without altering nuclear staining (Scale=30µm).

These results demonstrate that bafilomycin A1 effectively disrupts ACS-induced vacuolar acidification. Acridine orange accumulates in acidic vesicular compartments and emits orange–red fluorescence upon protonation, serving as a reliable indicator of vacuolarpH. The lack of such fluorescence in bafilomycin A1–treated cells is consistent with inhibition of vacuolar-type H⁺-ATPase (V-ATPase) activity, which is required for proton translocation and maintenance of acidic vacuolarpH[44]. The retention of nuclear green fluorescence in both control and treated cells confirms that bafilomycin A1 selectively affects vacuolar acidification without altering nuclear staining.

### 3.7 Pharmacological Modulation of ACS-Induced Vacuolation

Neutral Red (NR) staining was employed to quantitatively assess vacuolar accumulation in HeLa cells treated with activated calcium sulfate (ACS) in the presence of ten different drugs (Figure 9). HeLa cells were seeded in a 96-well plate and allowed to adhere overnight. For drug screening, columns 1–10 were assigned to individual drugs, each applied as a serial dilution across eight wells within the respective column. A fixed volume (30 µL) of freshly prepared 5-min ACS suspension was added uniformly to all wells in columns 1–10.

**Figure 9.**
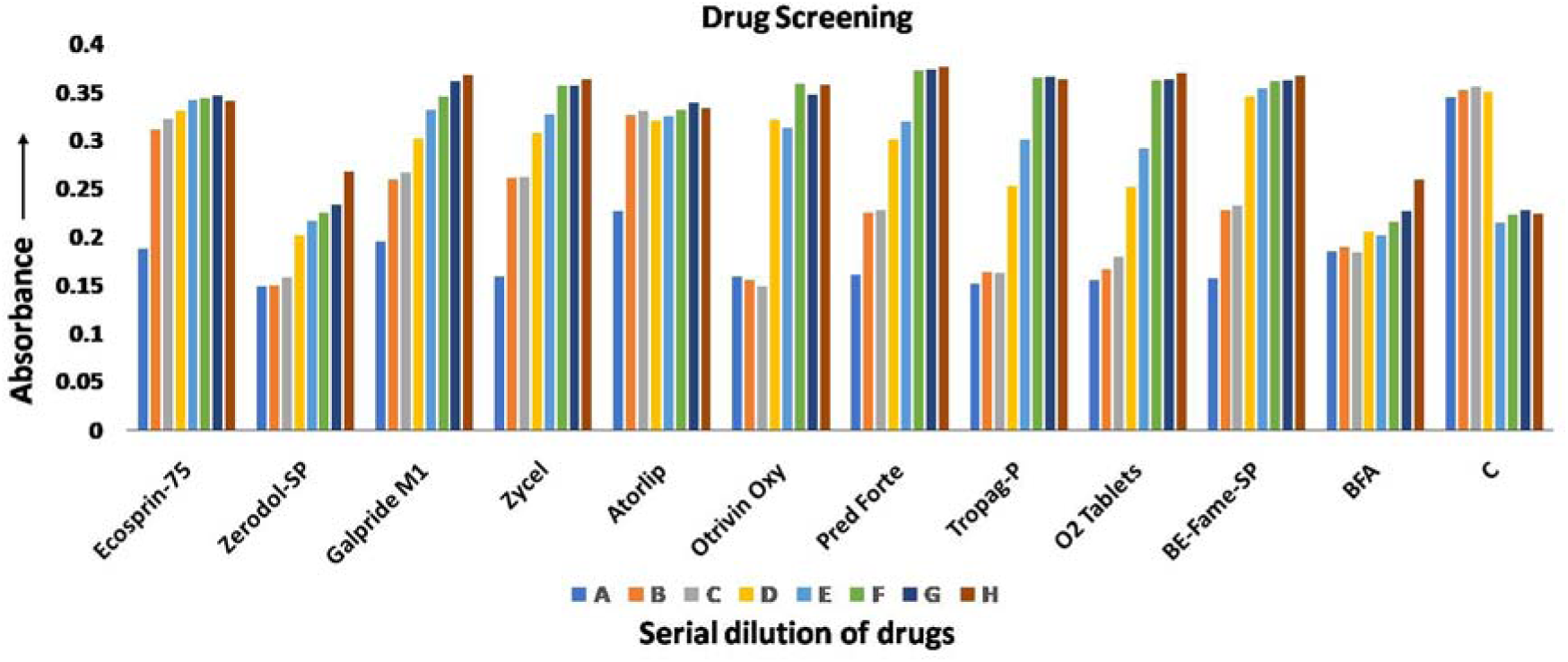
Neutral Red (NR)–based quantification of vacuole formation in ACS-treated HeLa cells under drug screening conditions. HeLa cells were treated with activated calcium sulfate (ACS) in the presence or absence of 11 drugs across a serial dilution (concentrations 1–8). The positive control (ACS alone) displayed strong NR uptake, confirming robust vacuole induction, while the negative control (untreated cells) exhibited minimal NR staining, representing basal lysosomal activity. Drug-specific responses were observed: Drug Ecosprin-75, Atorlip, and Otrivin Oxy showed no inhibition, maintaining NR uptake similar to ACS control; Zerodol-Sp caused marked suppression of NR uptake at low concentrations, indicative of cytotoxicity, with partial recovery at higher doses; Galpride-M1(500mg) showed graded inhibition at the lowest concentration followed by progressive restoration of vacuole formation; Drugs Pred Forte, Torpag-P, O2 and BE-Fame-sp induced cytotoxicity at higher concentrations with subsequent recovery at lower doses, as evidenced by restored NR accumulation. Data highlight the ability of ACS-induced vacuolation, quantified by NR staining, to discriminate between cytotoxic suppression and direct inhibition of vacuole biogenesis, establishing this assay as a functional platform for drug screening.

**Figure 10.**
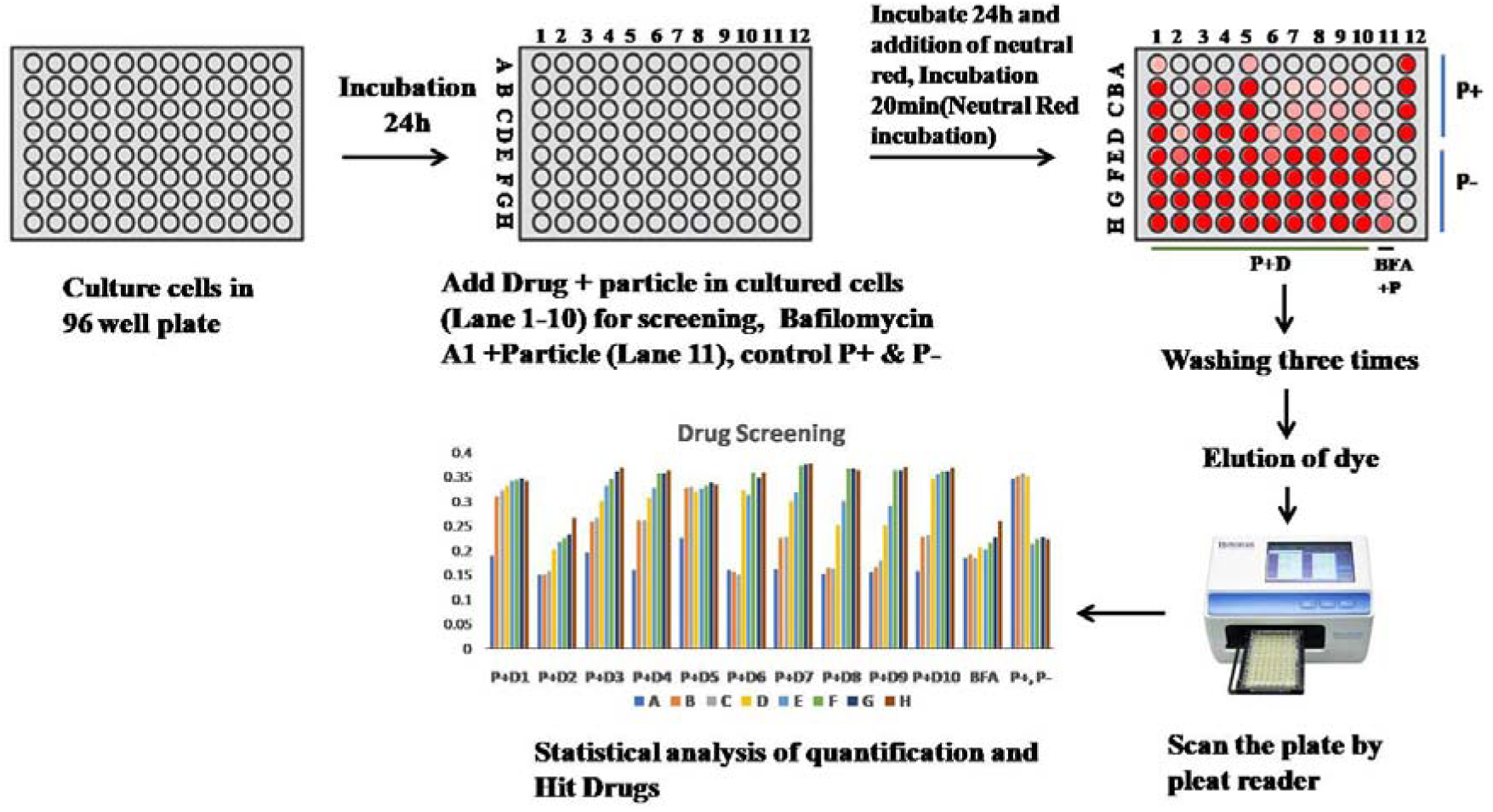
High-throughput drug screening based on vacuole quantification in ACS-treated cells. Schematic representation of the experimental workflow used for pharmacological screening of compounds that modulate activated calcium sulfate (ACS)–induced vacuolation. Cells were first seeded in a **96-well microplate** and allowed to adhere under standard culture conditions. Cells were treated with different drugs in lanes 1–10, followed by exposure to activated calcium sulfate (ACS) particles to induce vacuole formation. Lane 11 received Bafilomycin A1 (BFA1) prior to ACS treatment and served as the positive control for vacuole inhibition. Lane 12 served as the particle control, where the first four wells received ACS particles while the remaining four wells were left untreated. Vacuole formation was subsequently quantified using a plate-based assay, and absorbance values were measured using a microplate reader. The resulting data were analyzed to compare the effects of different compounds on ACS-induced vacuolation.

Column 11 served as the drug-positive control, where the first four wells received 51.35, 48.57, 45.45 and 41.9nM (high concentration) of Bafilomycin A1 (BFA) working solution(100nM), respectively, and the remaining four wells received 28, 21.74, 14.29, and 5.26nM (low concentration) of BFA. Column 12 served as the control column, with the first four wells receiving ACS alone (without drugs) and the remaining wells containing untreated cells without ACS or drugs. After 24 hrs of culture, ACS-treated control cells exhibited robust NR uptake, confirming consistent vacuole induction by ACS, whereas untreated cells displayed minimal NR accumulation, reflecting basal lysosomal activity (Figure 9).

Distinct drug-specific response patterns were observed across the concentration gradient (wells 1–8). Drug 1 showed no inhibitory effect on vacuole formation, with NR uptake closely matching ACS-treated controls and exceeding the lowest concentration of the drug-positive BFA control, indicating that vacuole biogenesis remained unaffected (Figure 9). In contrast, Drug 2 exhibited strong inhibition at lower concentrations, with the first three concentrations yielding NR values lower than the untreated control and below the highest concentration of the BFA control, indicative of pronounced drug-induced cytotoxicity (Figure 9). From the fourth concentration onward, NR uptake gradually increased and approached the levels observed in untreated and high-dose Bafilomycin A1 control cells, indicating recovery of cell viability and partial restoration of vacuole formation (Figure 9).

Drug 3 and 4 demonstrated transient inhibition at the first concentration, followed by dose-dependent recovery, with vacuole formation restored from the second concentration onward and ultimately reaching levels comparable to ACS-treated controls (Figure 9). Drug 5 showed no measurable inhibition of vacuolation, maintaining robust NR uptake similar to ACS-treated cells and exceeding the lowest concentration of the BFA control (Figure 9).

Drug 6 induced marked cytotoxicity at high concentrations (1st–3rd), as reflected by reduced NR uptake, followed by recovery at lower concentrations (≥4th), where vacuole formation resumed, evidenced by a progressive increase in NR accumulation (Figure 9). A comparable trend was observed for Drug 7, wherein the first concentration caused cell death, the second and third concentrations supported cell viability with suppressed vacuolation, and from the fourth concentration onward, vacuole formation was restored (Figure 9). Drugs 8, 9, and 10 exhibited response patterns analogous to Drugs 6 and 7, indicating early concentration-dependent cytotoxicity followed by recovery and re-establishment of vacuolar activity at higher concentrations (Figure 9).

Collectively, these results underscore the dual utility of ACS-induced vacuolation as both a sensitive readout of particle–cell interactions and a functional in vitro platform for drug screening. Drugs 2, 6, 7, 8, 9, and 10 displayed concentration-dependent cytotoxic effects at lower doses, followed by recovery and modulation of vacuole formation at higher concentrations. In contrast, Drugs 1 and 5 exerted negligible effects on vacuolar dynamics, suggesting limited involvement in phagocytosis-associated pathways. Drug 3 and 4 exhibited a graded inhibition–recovery profile, highlighting its potential role as a modulator of vacuolar dynamics rather than a purely cytotoxic agent.

The ACS-induced vacuolation assay, coupled with Neutral Red (NR)–based quantification, establishes a robust and functionally informative in vitro platform for screening modulators of phagocytosis-associated vacuole biogenesis [42]. The reproducible formation of large, acidified vacuoles in ACS-treated cells generates a high-contrast, quantifiable readout that sensitively captures drug-induced perturbations in vacuolar dynamics across a concentration gradient [31]. This system effectively discriminates between compounds that exert direct inhibitory effects on vacuole formation and those that suppress NR accumulation secondary to cytotoxicity, as evidenced by distinct inhibition–recovery profiles observed among the tested drugs. Importantly, the graded responses—ranging from complete resistance to vacuole inhibition, transient suppression with recovery, and concentration-dependent cytotoxicity followed by restoration of vacuolar activity—underscore the assay’s ability to resolve mechanistically diverse drug actions within a single experimental framework. Unlike conventional phagocytosis assays that rely on fluorescent labelling, end-point imaging, or specialized instrumentation [46], the ACS–NR platform offers a simple, low-cost, and scalable approach compatible with 96-well formats, making it well suited for preliminary drug screening. By directly linking particle-induced vacuolation with cellular viability and lysosomal acidification, this platform provides a physiologically relevant functional readout for identifying candidate inhibitors or modulators of phagocytic and vacuolar pathways, thereby offering clear advantages for early-stage screening of compounds targeting phagocytosis-related processes.

## 4. CONCLUSION

Activated calcium sulfate (ACS) provides a biocompatible, chemically simple, and cost-effective alternative to conventional phagocytic particles and commercial assay kits for interrogating phagosome–lysosome dynamics. Unlike polymeric or opsonized particles that depend on receptor-specific uptake or immune priming, ACS induces robust vacuolization, enabling unbiased assessment of phagocytic pathways in professional and non-professional phagocytes. ACS-induced vacuoles undergo progressive maturation and acidification, permitting dynamic evaluation of lysosomal function using straightforward colorimetric readouts such as neutral red, without the requirement for specialized probes or instrumentation [47]. Building on these properties, the ACS-based drug screening assay described herein enables rapid, sensitive, and economical high-throughput identification of compounds that modulate ACS phagocytosis and vacuole formation, offering a practical advantage over imaging and flow cytometry–dependent approaches [48]. While the assay efficiently captures overall phagocytic activity, ACS represents an inorganic particle and does not fully recapitulate the biological complexity of phagocytic targets such as bacteria or yeast.To address this limitation, we are developing a second-generation hybrid particle system that integrates ACS with bacterial and yeast components, enabling comparative analysis of inorganic, bacterial, and fungal phagocytosis using the same simple neutral red and acridine orange (NR/AO)–based readout.

## 5. AUTHORSHIP CONTRIBUTION STATEMENT

**VG**: Methodology, Investigation, Formal analysis, Data curation, **AF:** Methodology, Investigation, Formal analysis, **GD**: Methodology, Investigation, analysis, Data curation, Conceptualization, Writing original draft, review & editing, **AK**: Methodology, Investigation, **SB**: Methodology, Investigation, Formal analysis **AG**: Methodology, Investigation, Formal analysis, **SD**: Methodology, Investigation, **AJ**: review & editing, Visualization, Validation, Methodology, Investigation, Funding acquisition, Formal analysis, Data curation, Conceptualization. **SKD**: Writing, review & editing, Supervision, Resources, Project administration, Methodology, Funding acquisition, Conceptualization.

## 6. AKNODLEGEMENT

This research was supported by Science & Engineering Research Board (SERB) Core Research Grant (CRG/2022/009045), Department of Science and Technology, Government of India.We thank Dr. Neeladrisingha Das, Department of Radiation Oncology, School of Medicine, Stanford University, Palo Alto, CA, USA, for performing the experiment in the supplementary video. We also thank Digilab Bio Analytical Instruments Pvt. Ltd., India, for assistance with microscopy experiments.

## 8. SUPPLEMENTARY FIGURES AND TABLES

**Supplemantry Figure 1.**
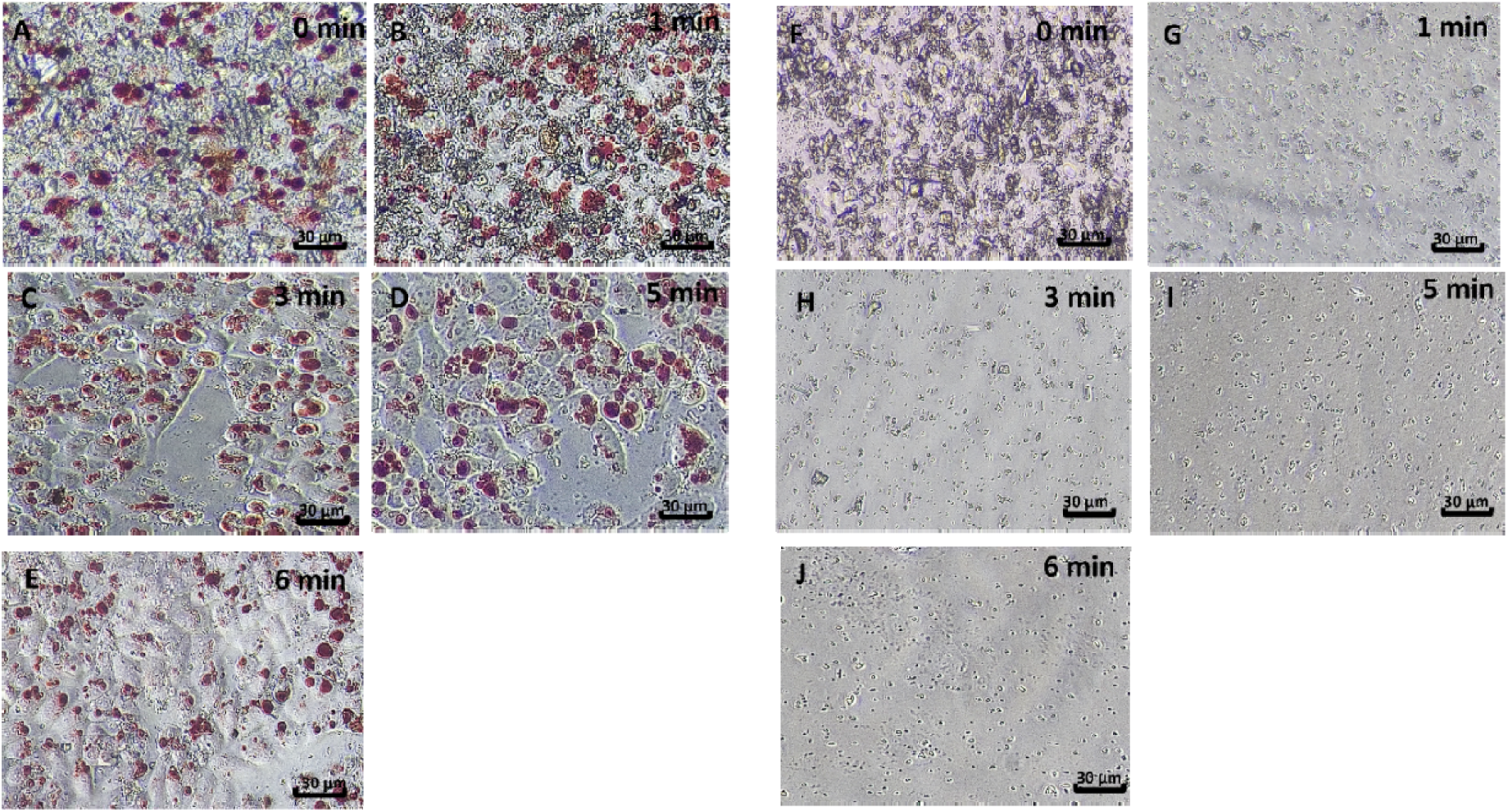
Particle heterogeneity and vacuolation during time-resolved sedimentation. Microscopic images of calcium sulfate suspensions collected at 1–3 minutes revealed highly heterogeneous particle populations with wide size variation and increased particle counts, corresponding to extensive vacuolation in HeLa cells. By the 4th–5th minute, suspensions transitioned toward homogeneity, with particle size distribution stabilizing and cell vacuolation appearing moderate and uniform across 5–9 minutes.

**Supplementary Figure 2.**
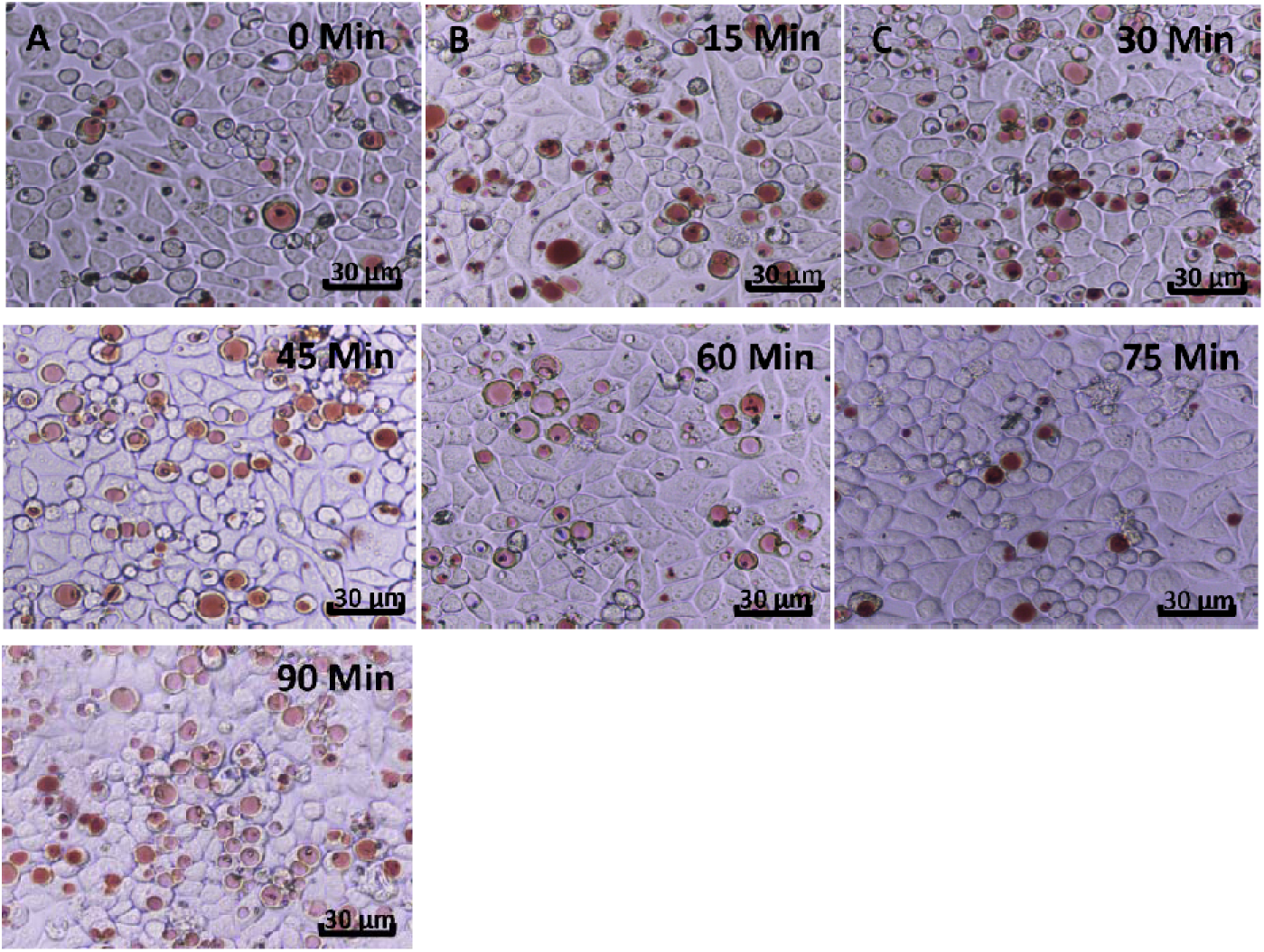
Immediate response of ACS-induced vacuoles to Bafilomycin A1. Microscopy images of HeLa cells with pre-formed vacuoles treated with Bafilomycin A1 and stained with neutral red (NR). Vacuoles actively took up NR at all observed time points (0–90 min), indicating that short-term exposure to Bafilomycin A1 did not impair vacuole acidification.

**Supplementary Figure 3.**
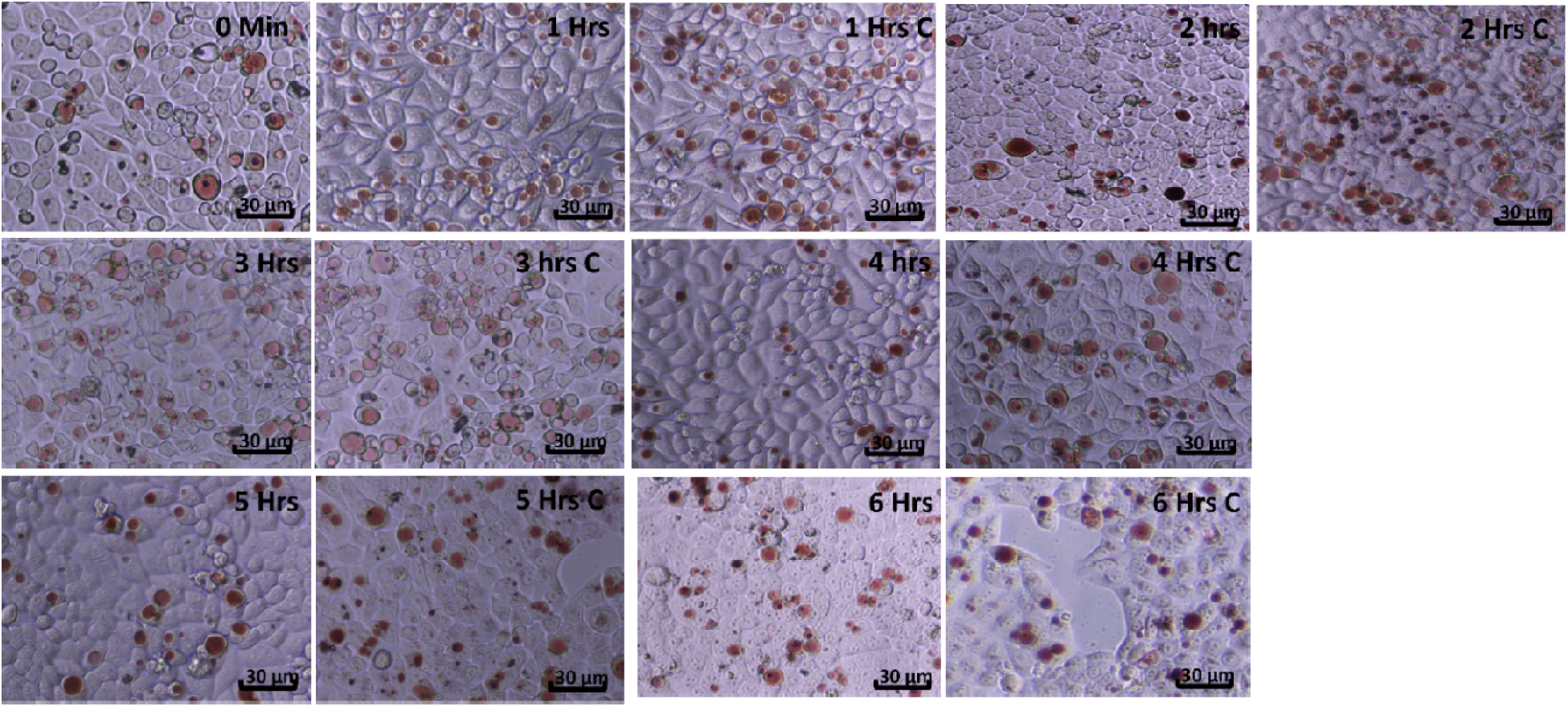
Hourly assessment of vacuole acidification following Bafilomycin A1 treatment. HeLa cells exposed to Bafilomycin A1 were monitored for 6 hours with NR staining at hourly intervals. Vacuoles continued to accumulate NR throughout the observation period, demonstrating that vacuole acidification remained intact during early exposure of Bafilomycin A1.

**Supplementary Figure 4.**
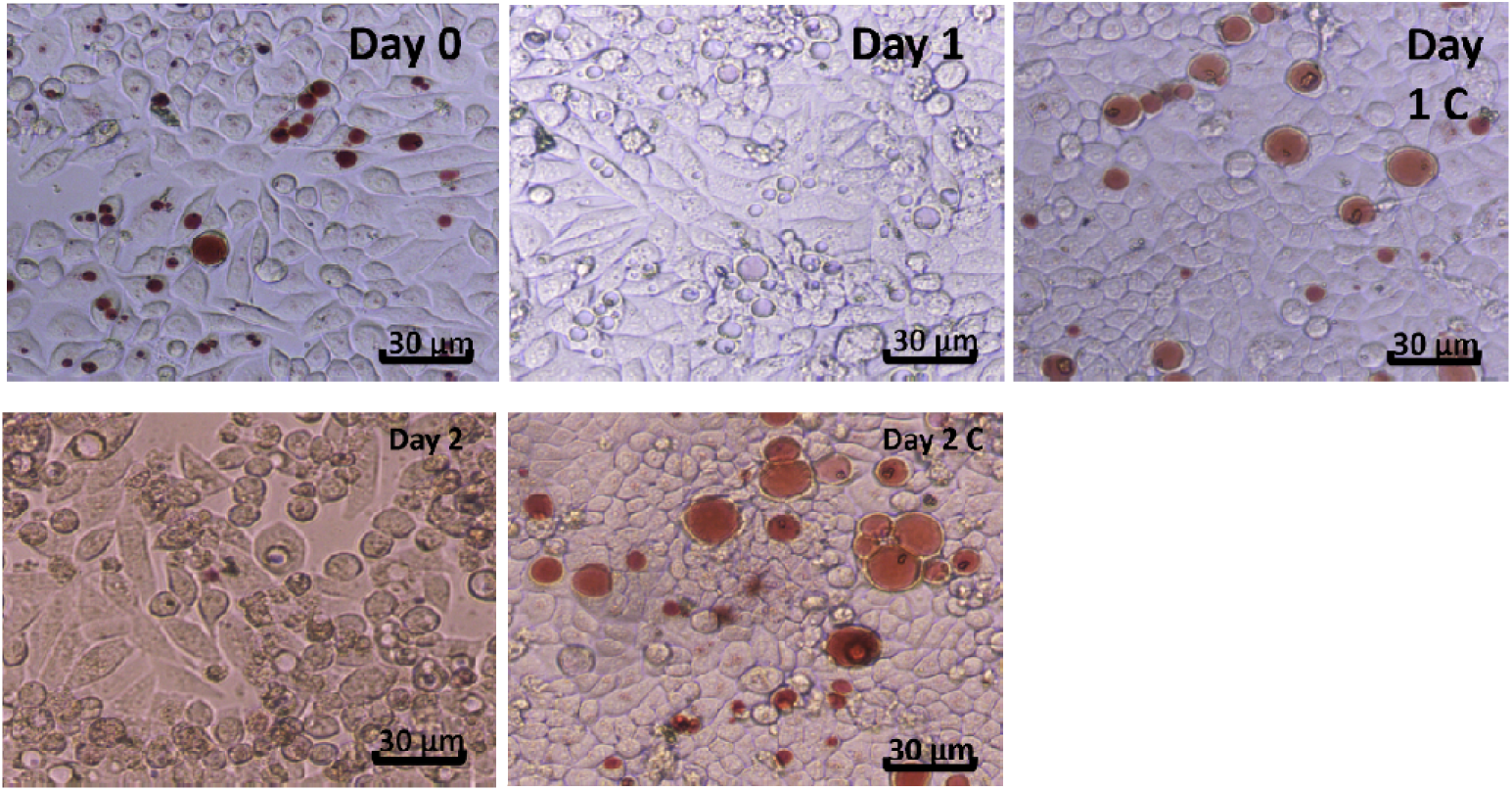
Delayed inhibition of vacuole acidification by Bafilomycin A1. Daily response analysis of HeLa cells treated with Bafilomycin A1 revealed NR-positive vacuoles on Day 0; however, vacuoles failed to take up NR after 24 hours (Day 1) and 48 hours (Day 2). These findings confirm that prolonged Bafilomycin A1 treatment abolishes vacuole acidification.

## 9. SUPPLEMENTARY VIDEO

**Time-lapse imaging of vacuole formation and turnover in A549 cells treated with 2-min stock sedimentation heterogeneous ACS particles.**

Time-lapse microscopy of A549 cells following treatment with 2-min stock sedimentation heterogeneous activated calcium sulfate (ACS) particles showed a progressive increase in vacuole number and size up to 24 h, indicating active particle internalization and vacuolar expansion. Beyond 24 h, both vacuole number and size gradually declined, reflecting intracellular degradation of ACS particles and vacuolar resolution. By day 4, a high degree of vacuolar turnover was evident; however, a few isolated vacuoles persisted. This persistence is consistent with the heterogeneous particle population in the 2-min sedimentation fraction, which contains larger ACS particles requiring extended time for complete intracellular degradation.

